# A Biologically Informed Heterogeneous Graph Neural Network for Multi-Task Prediction of ncRNA–Metastasis–Cancer Interactions

**DOI:** 10.64898/2026.08.18.745571

**Authors:** Farzad Midjani, Mohammadreza Shaghouzi, Amirhossein Dehqan Banadaki, Nima Rahimikashkooli, Fateme Zahra Keshtkar, Mahdi Malekpour, Samaneh Hashemi, Yasmin G. Hernandez-Barco, Saeed Soleymanjahi

**Affiliations:** Artificial Intelligence Clinical laboratory and Biological Data Bank, Shiraz University of Medical Sciences, Shiraz, Iran; Department of Medical Biotechnology, School of Advanced Medical Sciences and Technologies, Shiraz University of Medical Sciences, Shiraz, Iran; Systems Medicine Research Core, Shiraz University of Medical Sciences, Shiraz, Iran; Tarbiat Modares University, Tehran, Iran; Institute of Neuroscience and Medicine (INM-7: Brain and Behaviour), Research Centre Jülich, Jülich, Germany; Internal Medicine department, School of Medicine, Shiraz University Of Medical Sciences, Shiraz, Iran; Division of Gastroenterology, Mass General Brigham, Harvard School of Medicine, Boston, USA

**Author notes:** Corresponding author: Saeed Soleymanjahi.

## Abstract

Metastasis involves context-dependent molecular interactions in which non-coding RNAs, particularly miRNAs and circRNAs, play important regulatory roles. However, existing computational approaches generally do not jointly represent cancer type, metastatic event, and cancer-specific metastatic context. We developed a context-aware multi-task heterogeneous graph neural network (GNN) for predicting ncRNA associations with cancer types and metastatic events. The framework integrates multiple biological repositories into a heterogeneous graph representing ncRNAs, cancers, metastatic event types (METs), and cancer-specific metastatic instances (CSMIs). The model performs six link-prediction tasks using a hierarchical transformer-based encoder and multi-relational TuckER decoder. Across ten independently initialized runs evaluated on the RNA-group-disjoint held-out test set, the model achieved a global AUROC of 0.8801 ± 0.0118 and an F1 score of 0.8260 ± 0.0071. All three ablation variants yielded lower AUROC, with the largest reduction under independent task training. Case studies in pancreatic cancer, colorectal cancer, and hepatocellular carcinoma provided disease-level, event-level, and expression-based support, respectively, for top-ranked candidate associations. The framework enables context-specific prioritization of ncRNA–cancer–metastasis associations for experimental evaluation.

## 1 Introduction

Cancer remains a major threat to human health, encompassing a heterogeneous group of diseases characterized by uncontrolled cell proliferation (Sung et al., 2021). Prognosis worsens substantially once cancer spreads to distant sites, as disseminated tumor cells can survive, adapt to diverse tissue environments, and evade therapeutic intervention (Gerstberger et al., 2023). These capabilities promote treatment resistance and disease recurrence, making metastasis a major contributor to cancer-related mortality (Dillekås et al., 2019).

For decades, cancer research was largely guided by the central dogma of molecular biology, emphasizing the flow of information from DNA to RNA to protein and focusing primarily on mutations in protein-coding genes and dysregulated signaling pathways (Slack and Chinnaiyan, 2019). However, the Human Genome Project and next-generation sequencing revealed that much of the human genome is transcribed into non-coding RNAs (ncRNAs) (Veneziano et al., 2016). Although ncRNAs generally do not function through protein-coding capacity and were once dismissed as transcriptional “junk” or artifacts, they are now recognized as important regulators of cellular function and gene expression (Hentze et al., 2018; Walter, 2024). Their dysregulation is a common feature of cancer and can influence tumorigenesis and metastasis, with microRNAs (miRNAs) and circular RNAs (circRNAs) attracting particular interest because of their major roles in oncology (Kim et al., 2023).

ncRNA function is highly context-dependent: the same miRNA or circRNA may play different regulatory roles across cancer types and metastatic processes (Yarmishyn et al., 2022). Moreover, interactions such as the circRNA– miRNA–mRNA axis create dense regulatory networks in which local perturbations can influence broader metastatic behavior (Jia et al., 2024). Existing computational approaches have provided useful insights, but they often model ncRNA associations as isolated pairs or broad disease links rather than as cancer- and metastasis-aware regulatory networks (Hu et al., 2023). Because these interactions are inherently relational, heterogeneous GNNs are well suited to integrate ncRNAs, cancer types, metastatic processes, and their dependencies (Rosa et al., 2022).

To address these limitations, we developed a context-aware multi-task heterogeneous GNN that predicts six classes of ncRNA–cancer–metastasis links and explicitly represents cancer-specific metastatic instances through CSMI nodes. The study contributes: (i) a heterogeneous graph formulation that integrates ncRNAs, cancer types, metastatic events, and cancer–event-specific contexts; (ii) a hierarchical two-stage heterogeneous GNN with task-specific adaptation and TuckER-based scoring; and (iii) an evaluation framework combining RNA-group-disjoint held-out testing, seed-level stability analysis, component ablation, and external evidence assessment.

## 2 Methods

Figure 1 summarizes the data-integration, representation-learning, and link-prediction workflow.

**Figure 1:**
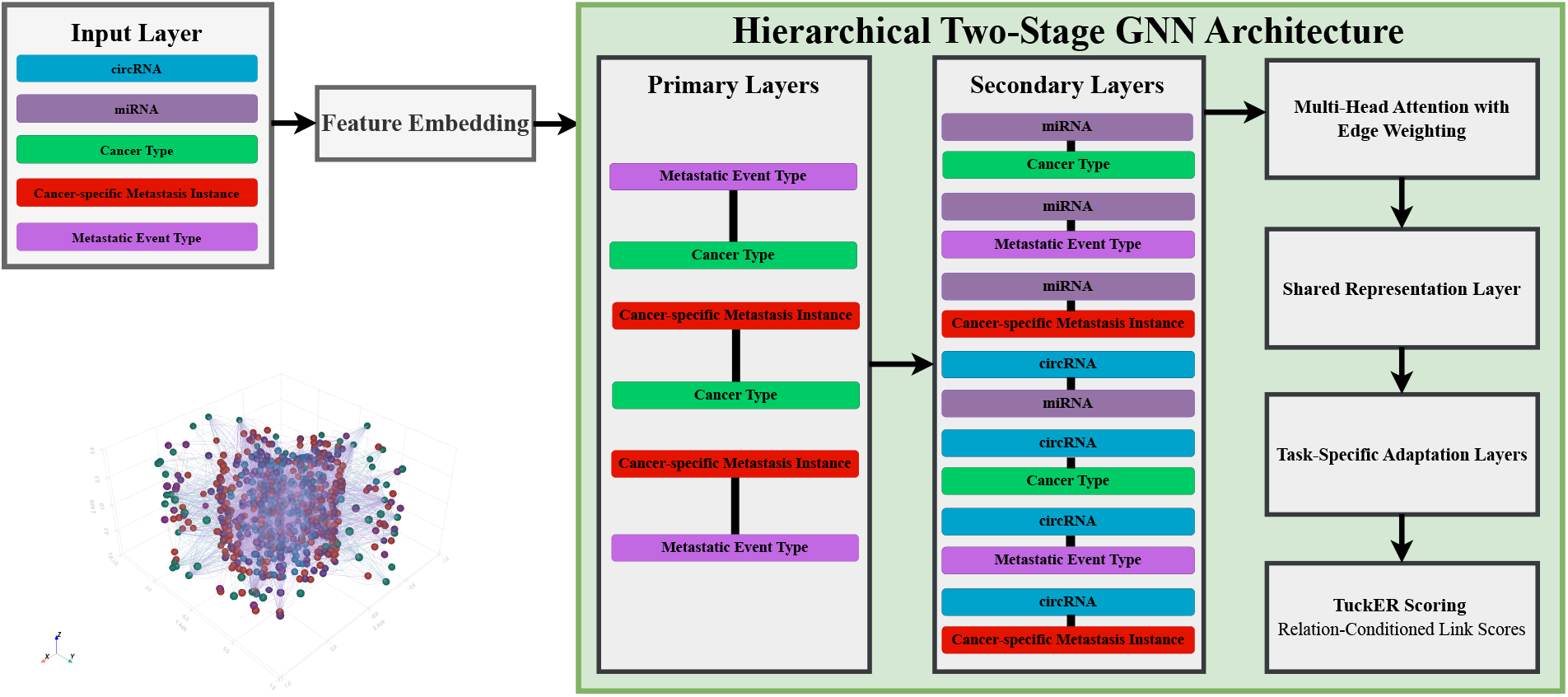
Overview of the framework. Empirical ncRNA features and trainable contextual-node embeddings are integrated within a heterogeneous graph. Sequential contextual and ncRNA-regulatory message passing stages is followed by shared representation learning, task-specific adaptation, and TuckER-style scoring across six link-prediction tasks.

### 2.1 Dataset Construction and Preprocessing

#### 2.1.1 Data Sources and Acquisition

We integrated four complementary biological repositories. ncR2Met provided experimentally supported or literature-curated associations between differentially expressed ncRNAs and cancer metastatic events across 32 metastatic processes and 53 human malignancies (Yu et al., 2023). CircBank supplied circRNA–miRNA interaction pairs, representing a key regulatory axis through which circRNAs can modulate miRNA activity (Liu et al., 2019; Salmena et al., 2011; Memczak et al., 2013; Hansen et al., 2013). dbDEMC provided cancer-specific miRNA differential-expression profiles based on logFC values, which were transformed into initial miRNA node features (Xu et al., 2022). CircAtlas supplied circRNA expression profiles based on read counts per million, which were used to initialize circRNA node features (Wu et al., 2020).

#### 2.1.2 Node and Edge Definition for the Heterogeneous Graph

#### 2.1.2.1 Node Types

The heterogeneous graph contains five node types: miRNA, circRNA, Cancer, MET, and CSMI. miRNA and circRNA nodes represent regulatory ncRNAs, Cancer nodes represent malignancy types, MET nodes represent metastatic processes, and each CSMI node represents an observed Cancer–MET combination, enabling prediction at cancer-level, event-level, and cancer–event-specific resolution (Figure 2).

**Figure 2:**
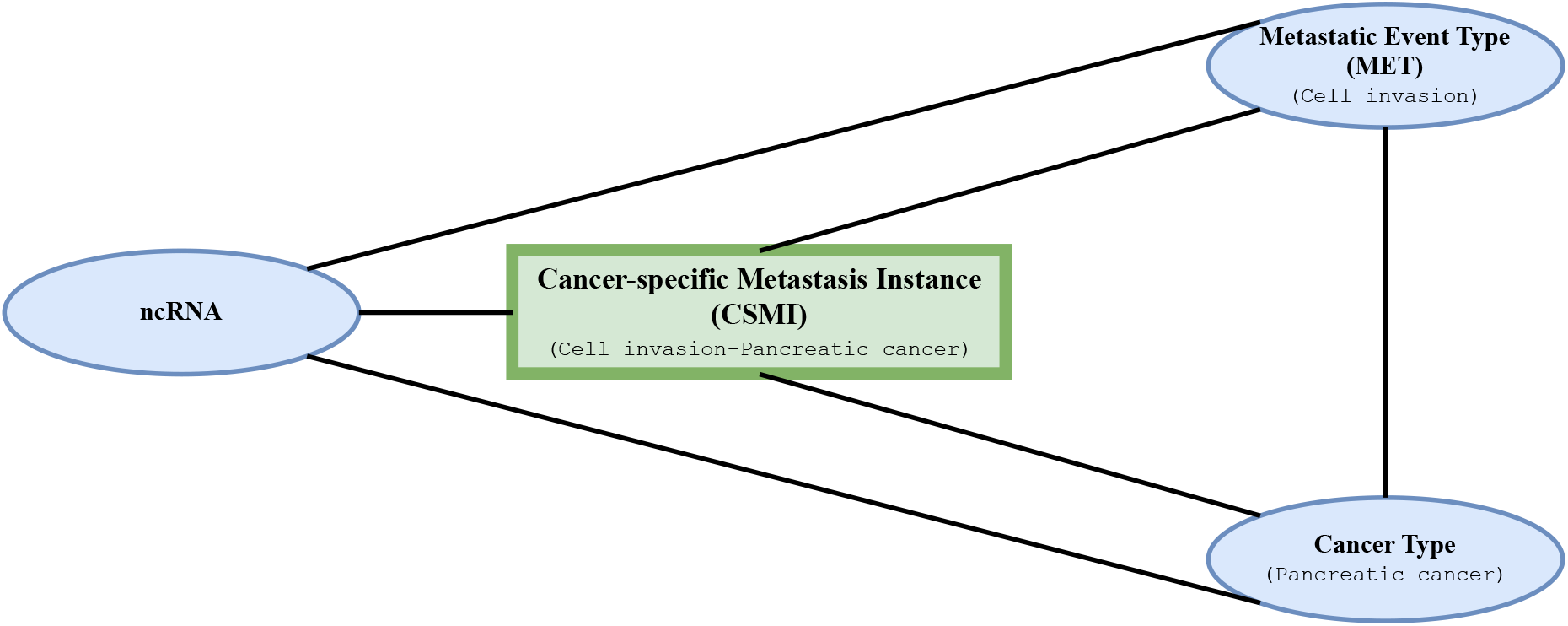
Construction of a cancer-specific metastatic instance. The MET node represents a general event, such as cancer cell invasion, and the Cancer node represents a malignancy, such as pancreatic cancer. Their paired context defines the CSMI node. Separate miRNA and circRNA relations connect ncRNAs to Cancer, MET, and CSMI targets.

#### 2.1.2.2 Edge Types

The graph contained ten directed forward relation types: seven ncRNA-related relations and three contextual relations connecting CSMI, MET, and Cancer nodes. After split-specific masking of target edges, reverse counterparts were added only for retained message-passing edges, yielding up to 20 directed message-passing relation types in each partition.

#### 2.1.3 Graph Construction and Feature Initialization

After identifier normalization, the five node sets and typed relation sets were assembled as a heterogeneous graph. Missing miRNA and circRNA feature values were retained during graph construction. For each model fit, missing values were replaced during the forward pass with node- and feature-specific trainable parameters initialized using Xavier uniform initialization and optimized jointly with the model. Observed values remained unchanged, preserving the original 80-dimensional miRNA and 13-dimensional circRNA feature representations.

For an RNA node *v* of type *τ*(*v*), the empirical feature vector **x**_*v*_ was projected into the 12-dimensional hidden space:

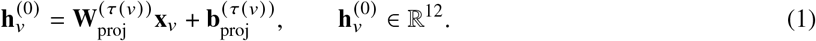

The projection parameters were learned separately for miRNA and circRNA nodes. Cancer, MET, and CSMI nodes lacked empirical input features and were initialized using separate trainable 12-dimensional embeddings with Xavier-uniform initialization (Glorot and Bengio, 2010).

Each relation was converted into a typed adjacency matrix. For miRNA–Cancer, miRNA–CSMI, circRNA–Cancer, and circRNA–CSMI relations, expression direction was encoded as an edge attribute: +1 for upregulation and −1 for downregulation.

#### 2.1.4 Data Splitting and Negative Sampling

Observed positive edges from the six prediction tasks were first divided into an 80% development partition and a 20% held-out test partition using a group-aware split in which the biological RNA source was the grouping variable. Consequently, each miRNA or circRNA source was assigned to only one outer partition across all six tasks. Within the development partition, four-fold stratified group cross-validation was performed, with prediction task used for stratification and RNA source used for grouping. Ten percent of each fold’s fitting-positive groups was reserved through a second group-aware split for validation, checkpoint selection, early stopping, and threshold selection.

Within each fitting partition and prediction task, 80% of fitting positive edges were retained as target-relation message-passing edges and excluded from positive loss supervision. The remaining 20% were removed from the target-relation topology and used as positive supervision for the link-prediction loss. This split was performed after the outer and inner RNA-group partitions were fixed. Validation and held-out test positives were excluded from the encoder topology. Non-target contextual relations remained available for message passing in every partition, and reverse message-passing relations were created only after target-edge masks had been applied.

Negative examples were generated separately for each data partition and prediction task after the positive-edge split. Candidate pairs were restricted to source nodes represented in the current partition, while all observed positive pairs and negative pairs previously reserved for other partitions were excluded. Each task used a 1:1 positive-to-negative ratio, subject to the availability of eligible absent pairs. Negative candidates were absent typed source–destination pairs after exclusion of every observed positive pair and every pair reserved for another partition. Candidates were grouped by shortest-path distance in an undirected view of the fitting-only message-passing graph. Sampling targets were 30% from distance five, 30% from distance four, 30% from distance three, and 10% from the remaining eligible absent-pair pool. When a distance bucket contained too few candidates, its deficit was transferred to the next bucket.

### 2.2 Model Architecture

#### 2.2.1 The Hierarchical GNN Encoder

The encoder contains four sequential message-passing layers: two primary contextual layers followed by two secondary ncRNA-regulatory layers. Each layer uses a separate graph transformer convolution for each active directed relation in that stage. To stabilize training and mitigate over-smoothing and vanishing gradients (Oono and Suzuki, 2020), we apply layer normalization (Ba et al., 2016), LeakyReLU activation, dropout (Srivastava et al., 2014), and residual connections (He et al., 2016) after each layer.

Initial RNA features and contextual-node embeddings were projected into a shared 12-dimensional encoder space, as described in Section 2.1.3.

#### 2.2.1.1 Convolution Building Block

After partition-specific masking of target edges, reverse relations were created for retained message-passing edges. Each active directed relation was processed by a separate graph transformer convolution module. For a target node *i* and its neighboring nodes *j* ∈*N*(*i*) connected by a relation *r*, the update is driven by a multi-head attention mechanism. The output of a single attention head *k* is calculated as:

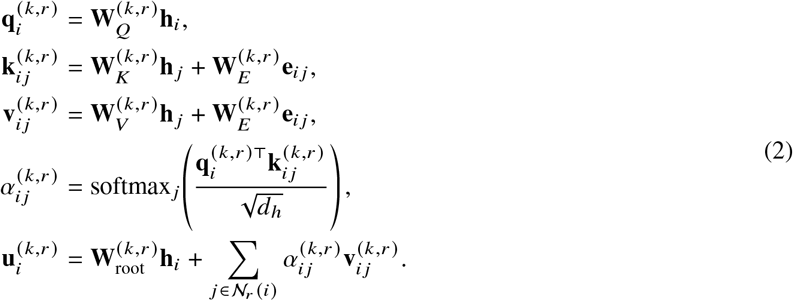

Here, 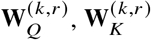 and 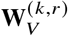 are relation- and head-specific projection matrices, and 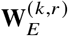 transforms the scalar edge attribute. Each relation-specific module used two 12-channel attention heads, producing a 24-dimensional concatenated representation. For each destination-node type, learned scalar relation weights were normalized across the active incoming relations and rescaled by the number of active relations:

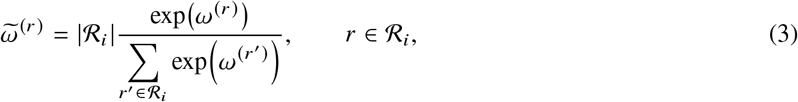

where *ω*^(*r*)^ is the learned scalar logit for relation *r*, and *R*_*i*_ is the set of active incoming relations for the destination-node type of node *i*. The weighted relation-specific outputs were then summed and projected to 12 dimensions before normalization, activation, dropout, and residual addition:

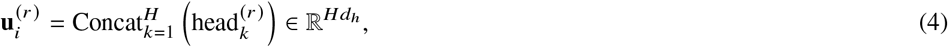

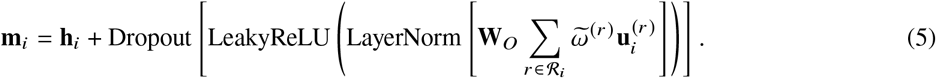

where 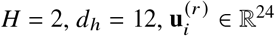 and **W**_*o*_ ∈ R^12×24^.

##### 2.2.1.2 Hierarchical Architecture: Primary and Secondary Passes

The encoder partitions the ten forward relation families and their retained reverse counterparts into two sequential message-passing stages (Figure 3). The primary stage processes the three context-defining forward relations CSMI →MET, CSMI →Cancer, and MET→ Cancer together with their retained reverse counterparts. The secondary stage processes the seven ncRNA-related forward relations and their retained reverse counterparts, connecting miRNAs and circRNAs to CSMI, MET, Cancer, and miRNA nodes. Relation-specific outputs are weighted and aggregated within each stage.

**Figure 3:**
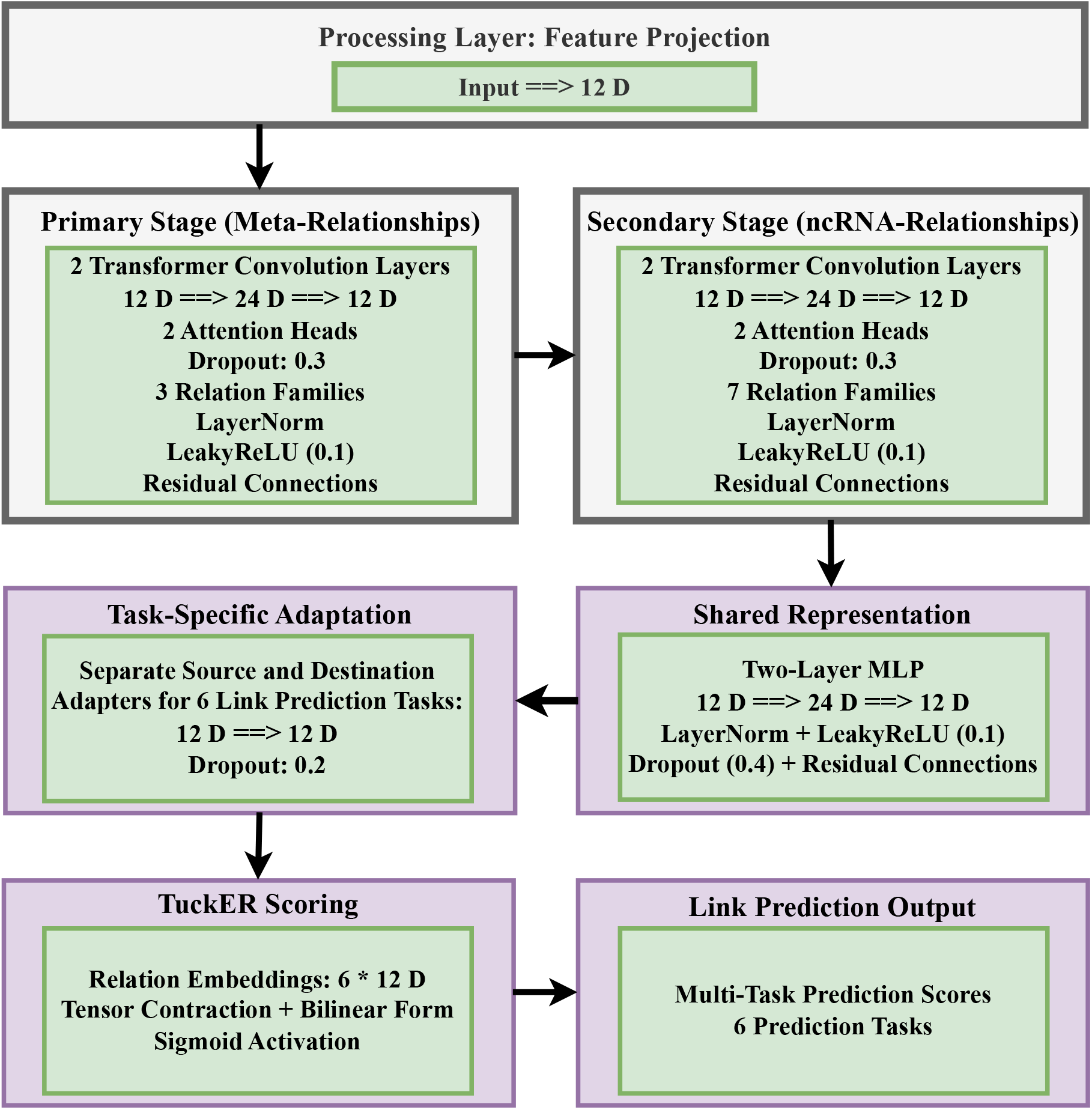
Sequential architecture of the multi-task link-prediction model. Variable-dimensional ncRNA features and trainable contextual-node embeddings are projected into a shared 12-dimensional space. Two primary graph transformer convolution layers process contextual relations, two secondary graph transformer convolution layers process ncRNA-related relations, and 12-dimensional task-specific adapters feed a TuckER-style decoder for six link-prediction tasks.

For stage *s* ∈ {primary, secondary}, the resulting aggregate is denoted by **g**^(*l*)^, and node *i* is updated as

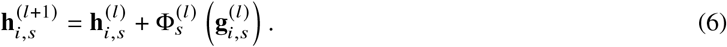

Here, 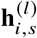is the previous-layer embedding for node *i* in stage 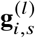 is the relation-weighted aggregate message for that stage, and 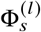 denotes the stage-specific projection, normalization, activation, and dropout block. The residual term preserves the previous-layer representation, while the sequential primary and secondary stages integrate contextual structure before ncRNA-related regulatory information.

##### 2.2.1.3 Global Residual Connection and Final Encoder Output

After the two primary and two secondary message-passing layers, the encoder added the initial projected embedding to the final secondary-stage representation to preserve node-specific information across the stacked layers (Oono and Suzuki, 2020; He et al., 2016):

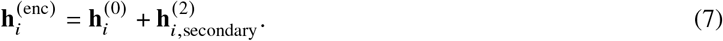

#### 2.2.2 Shared Representation Layer

The encoder output for each node type was refined by a type-specific residual MLP with dimensions 12 →24→ 12, LayerNorm, LeakyReLU activation, and dropout. The resulting shared embedding, 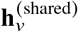, is defined as:

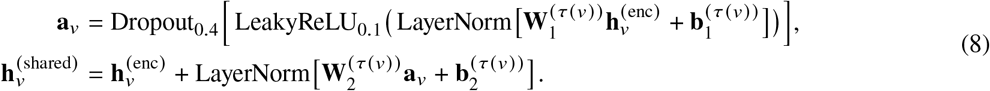

Here, (**W**_1_, **b**_1_) and (**W**_2_, **b**_2_) are the learnable parameters of the two MLP layers for node type *τ*(*v*).

#### 2.2.3 Task-Specific Adaptation Layers

For each prediction task *T*, separate source and destination feed-forward adapters transformed the shared embeddings into task-specific representations:

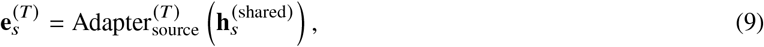

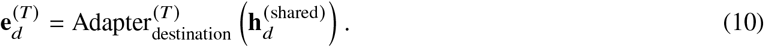

Each task used independent 12-dimensional source and destination adapters comprising a linear layer, LayerNorm, LeakyReLU activation, and dropout of 0.2. These adapters transformed the shared node representations separately for the six prediction relations.

#### 2.2.4 The Multi-Task Link Prediction Decoder: TuckER Scorer

The decoder applied a task-specific TuckER scoring function to the adapted source and destination embeddings for each of the six prediction relations (Balazevic et al., 2019). For task *T*, the decoder assigned a logit to triplet (*h*,T,*t*)using a TuckER-style bilinear function:

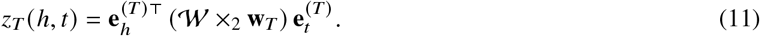

The corresponding bounded interaction score was

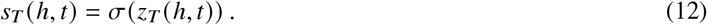

Here, 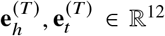 are the task-adapted source and destination embeddings, **w**_*T*_ ∈ R^12^ is the task relation embedding, and *W* ∈ R^12×12×12^ is the shared TuckER core tensor.

#### 2.2.5 Multi-Task Training Process

For a mini-batch *B*, losses were computed separately for the set of tasks represented in that batch, *T*_*B*_, and then averaged:

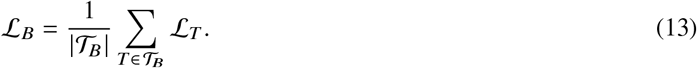

For task *T*, binary cross-entropy with logits was used. When the sampled fitting set was not exactly balanced because of limited eligible negatives, a task-specific positive-class weight was computed from the fitting examples as

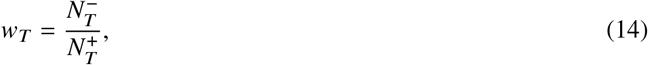

where 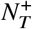 and 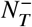 denote the numbers of positive and negative fitting examples for task *T*, respectively. The task-specific loss was

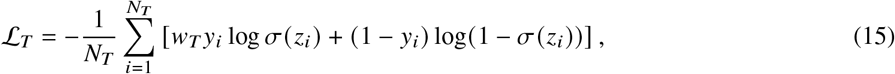

where *N*_*T*_ is the number of examples for task *T* in the mini-batch, *z*_*i*_ is the TuckER logit, and *y*_*i*_ ∈ {0, 1}.

#### 2.2.6 Evaluation Protocol

Cross-validation characterized development-set performance. In each final run, the validation partition was used only for model-state selection and global-threshold determination. The validation-selected model state and threshold were then applied once to the fixed RNA-group-disjoint held-out test partition.

Model performance was reported using AUROC, accuracy, precision, recall, and F1 score on the sampled evaluation sets. AUROC was used as the threshold-independent discrimination metric, while accuracy, precision, recall, and F1 score were computed using the validation-selected global threshold. A single global decision threshold was selected from the validation predictions by maximizing global F1 score and was then fixed before held-out test evaluation and all per-task threshold-dependent analyses. Downstream hypothesis generation ranked candidate associations using continuous model scores. Test labels were not used for model-state selection or threshold tuning. For stability assessment, the final training and held-out evaluation process was repeated across ten independently initialized runs, and the held-out metrics are reported as mean ± sample standard deviation.

### 2.3 Ablation Study Design

To assess multi-task sharing, contextual encoding, and graph topology, we evaluated three ablation variants across ten independently initialized runs using the same RNA-group-disjoint outer partitions as the full model.

The evaluated variants were: (i) *Independent task training*, in which one model was trained for each prediction relation and no encoder, representation, adapter, or decoder parameters were shared across tasks; these models used task-specific validation thresholds for threshold-dependent metrics; (ii) *Primary context backbone knockout*, in which the primary CSMI–MET–Cancer message-passing stage was removed while the secondary ncRNA-relational stage and task decoders were retained; and (iii) *Message-topology randomization*, in which message-passing edges were randomized independently within each relation type while preserving relation-specific edge counts, with all other training and evaluation settings unchanged.

### 2.4 Implementation Details

The model used 12-dimensional hidden representations, two attention heads, and 12-dimensional TuckER relation embeddings with a 12 × 12 × 12 core tensor. Dropout rates were 0.3 in the graph transformer convolution modules, 0.4 after message aggregation and in the shared MLPs, and 0.2 in the task-specific adapters. All models used AdamW with an initial learning rate of 0.001, weight decay of 0.001, and gradient clipping at 1.0 (Loshchilov and Hutter, 2019). Cross-validation models used a fixed learning rate. During final model fitting, ReduceLROnPlateau was additionally applied in maximization mode with a reduction factor of 0.5 and patience of five epochs, using validation AUROC as the monitored metric. Models were trained for up to 150 epochs with a batch size of 64 and early-stopping patience of 10 epochs. Data preprocessing used Pandas (McKinney, 2010) and NumPy (van der Walt et al., 2011), model implementation used PyTorch and PyTorch Geometric (Paszke et al., 2019; Fey and Lenssen, 2019; Fey et al., 2025), and evaluation used scikit-learn (Pedregosa et al., 2011).

### 2.5 Case Study Design and External Evidence Assessment

We assessed top-ranked candidate associations absent from the initial dataset through three external evidence analyses. For each case-study task, all eligible unobserved pairs were scored using the validation-selected model. Candidates were ranked by predicted probability, and pairs present in the training, validation, or held-out positive-edge sets were excluded before selection. For pancreatic cancer, we evaluated the top ten miRNA–cancer predictions using RNADisease v4.0 as disease-level external support (Chen et al., 2023). For colorectal cancer (CRC), we evaluated the top fifteen miRNA–CSMI and top fifteen circRNA–CSMI predictions through targeted literature review. Support was assigned when direct experimental evidence in CRC matched the predicted metastatic event or phenotype. Predictions without matching evidence in the targeted search were retained as model-prioritized candidates.

For the hepatocellular carcinoma (HCC) expression-based case study, the top 15 miRNA–CSMI predictions for the cancer cell migration–hepatocellular carcinoma context were evaluated using GEO Series GSE67139 (NCBI Gene Expression Omnibus, 2015; Xie et al., 2016). This dataset contains non-coding RNA array profiles from human HCC tumors with and without vascular invasion. Vascular-invasion-positive (VI-positive) tumors were compared against vascular-invasion-negative (VI-negative) tumors, while HCC cell-line samples were excluded from the analysis. Differential expression was assessed using GEO2R (NCBI Gene Expression Omnibus), which performs group-wise expression comparison on GEO datasets using the supplied processed data. Candidate miRNAs were matched to GEO2R probe identifiers and summarized by predicted probability, probe mappability, logFC direction, and adjusted P-value. Positive logFC values indicate higher expression in vascular-invasion-positive HCC tumors, whereas negative logFC values indicate lower expression in vascular-invasion-positive HCC tumors.

### 2.6 Global Prediction-Landscape Analysis

To characterize predictions beyond the three case-study cancers, cancer types were ranked by the number of retained ncRNA–CSMI candidate associations above the ensemble threshold. The ten highest-ranked cancer types were selected. Within each selected cancer type, candidates were stratified by metastatic event, which defines the corresponding CSMI. ncRNAs were then ranked by the total number of retained ncRNA–CSMI associations and by the number of selected cancer types in which they occurred.

## 3 Results

### 3.1 Summary of the Constructed Biological Graph

The constructed heterogeneous graph contained 807 nodes across five node types: 315 miRNAs, 193 circRNAs, 53 Cancer nodes, 32 MET nodes, and 214 CSMI nodes. It contained 5,267 unique directed forward edges across ten relation types, including ncRNA–CSMI, ncRNA–Cancer, ncRNA–MET, circRNA–miRNA, and contextual CSMI–Cancer/MET relations. Reverse counterparts were added only to partition-specific message-passing graphs and are not included in these forward-edge totals.

### 3.2 Cross-Validation Performance

Table 1 reports four-fold cross-validation performance within the 80% development partition.

**Table 1.** Four-fold stratified group cross-validation performance within the 80% development partition using validation-selected model checkpoints.

| Fold | AUROC | Accuracy | F1 Score | Precision | Recall |
| --- | --- | --- | --- | --- | --- |
| 1 | 0.87 ± 0.02 | 0.81 ± 0.02 | 0.82 ± 0.01 | 0.79 ± 0.03 | 0.85 ± 0.03 |
| 2 | 0.85 ± 0.02 | 0.80 ± 0.02 | 0.81 ± 0.02 | 0.76 ± 0.02 | 0.87 ± 0.02 |
| 3 | 0.87 ± 0.02 | 0.81 ± 0.01 | 0.82 ± 0.01 | 0.78 ± 0.03 | 0.86 ± 0.04 |
| 4 | 0.87 ± 0.02 | 0.80 ± 0.02 | 0.81 ± 0.03 | 0.77 ± 0.03 | 0.86 ± 0.05 |
| Mean | 0.86 ± 0.02 | 0.81 ± 0.02 | 0.82 ± 0.02 | 0.78 ± 0.03 | 0.86 ± 0.04 |

Values are mean ± SD across ten independently initialized runs.

Across the ten independently initialized final-model runs, validation loss decreased from 0.6914 ± 0.0076 at the first epoch to 0.4531 ±0.0626 at the selected model checkpoints. These results show that optimization consistently reduced validation loss across runs.

Across all four cross-validation folds, median validation AUROC increased during the initial training epochs and subsequently stabilized (Figure 4).

**Figure 4:**
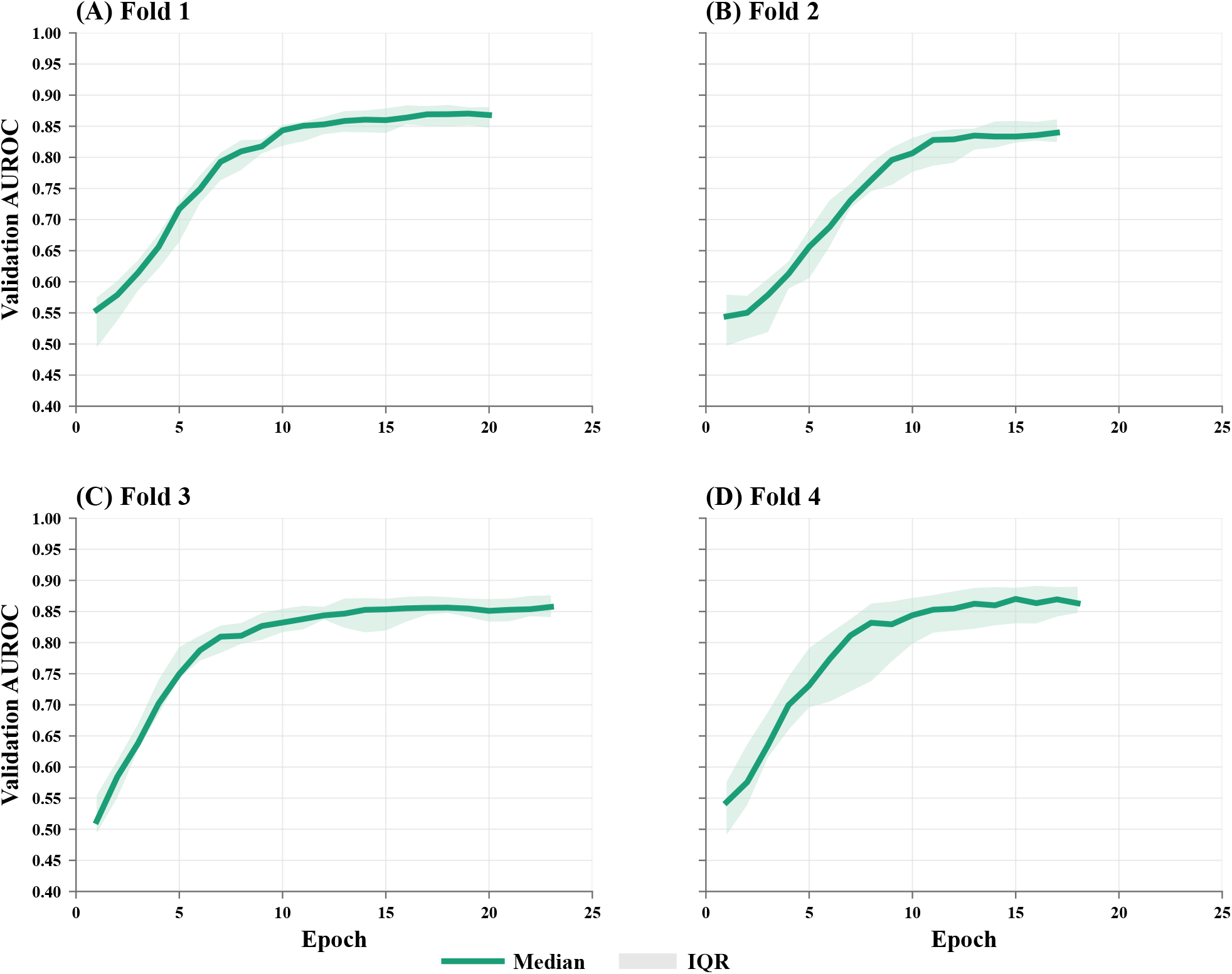
Validation AUROC trajectories across the four cross-validation folds. Lines show the median and shaded bands show the interquartile range across ten independently initialized runs.

### 3.3 Held-Out Test-Set Performance

Across ten independently initialized held-out runs, the validation-selected global threshold was 0.5250 ± 0.1080. On balanced sampled held-out test sets, the model achieved AUROC 0.8801 ± 0.0118, F1 score 0.8260 ± 0.0071, accuracy 0.8206 ± 0.0109, precision 0.8032 ± 0.0257, and recall 0.8513 ± 0.0213. Consistent with these results, the selected models achieved a test loss of 0.4573 ± 0.0250, closely matching the checkpoint validation loss of 0.4531 ± 0.0626. The close agreement between checkpoint validation and held-out test losses indicates little degradation on unseen test associations and is consistent with stable generalization within the evaluated RNA-group-disjoint setting.

Task-specific held-out metrics are reported in Table 2 and visualized in Figure 5. Figure 6 relates each task’s candidate-space burden to its held-out AUROC shortfall.

**Table 2:** Per-task held-out test performance across ten independently initialized runs. Values are mean ± SD.

| Interaction Type | Accuracy | F1 Score | Precision | Recall | AUROC |
| --- | --- | --- | --- | --- | --- |
| miRNA to CSMI | $0.74 \pm 0.03$ | $0.74 \pm 0.03$ | $0.74 \pm 0.04$ | $0.75 \pm 0.06$ | $0.78 \pm 0.04$ |
| circRNA to CSMI | $0.73 \pm 0.04$ | $0.74 \pm 0.04$ | $0.70 \pm 0.03$ | $0.80 \pm 0.08$ | $0.79 \pm 0.04$ |
| miRNA to Cancer | $0.81 \pm 0.04$ | $0.82 \pm 0.03$ | $0.78 \pm 0.05$ | $0.87 \pm 0.04$ | $0.85 \pm 0.02$ |
| circRNA to Cancer | $0.78 \pm 0.04$ | $0.79 \pm 0.05$ | $0.74 \pm 0.03$ | $0.86 \pm 0.10$ | $0.85 \pm 0.04$ |
| miRNA to MET | $0.93 \pm 0.02$ | $0.93 \pm 0.02$ | $0.93 \pm 0.03$ | $0.93 \pm 0.02$ | $0.95 \pm 0.02$ |
| circRNA to MET | $0.93 \pm 0.02$ | $0.93 \pm 0.02$ | $0.92 \pm 0.03$ | $0.94 \pm 0.02$ | $0.95 \pm 0.01$ |

**Figure 5:**
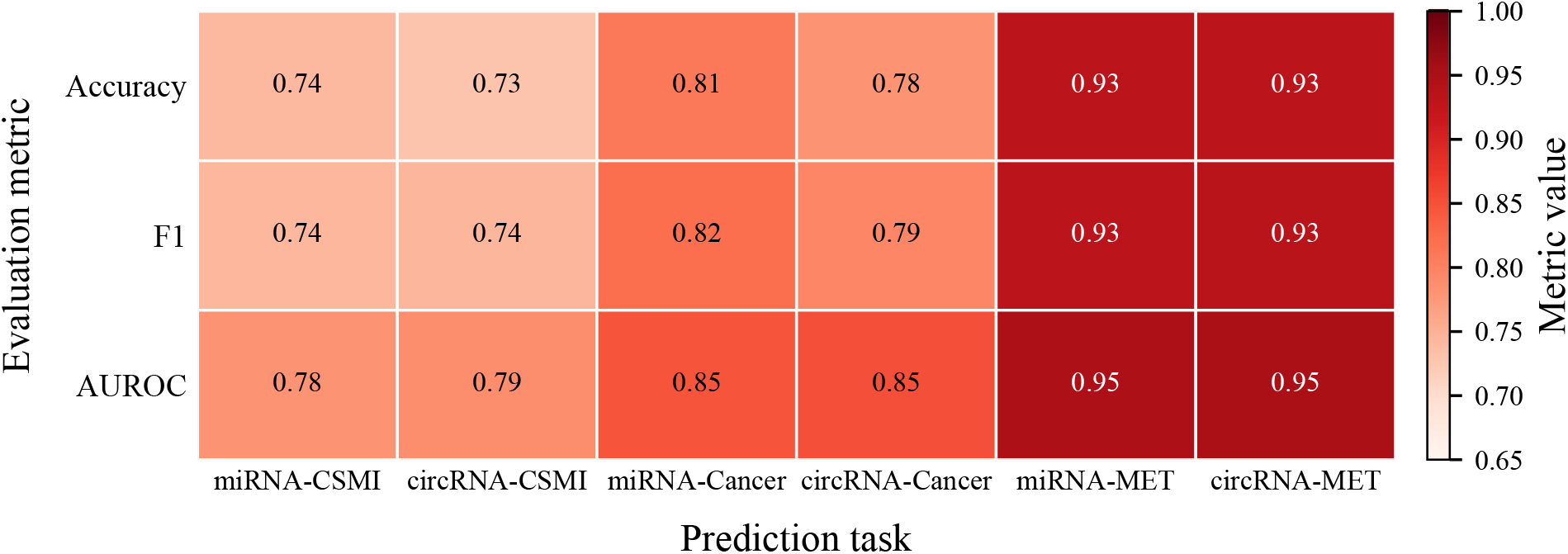
Task-wise held-out performance across the six link-prediction tasks. Cells report mean accuracy, F1 score, and AUROC across ten independently initialized runs. Corresponding sample standard deviations are reported in Table 2.

**Figure 6:**
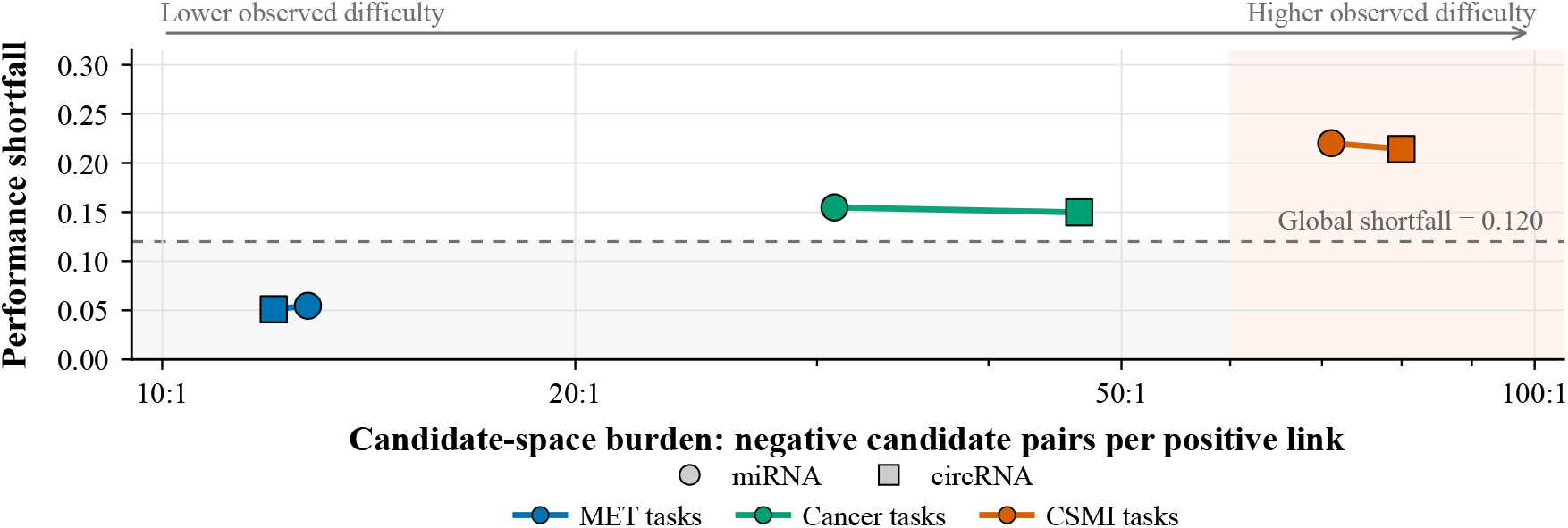
Task-difficulty landscape across the six ncRNA–metastasis–cancer link-prediction tasks. The x-axis shows candidate-space burden, defined as the ratio of possible negative candidate pairs to observed positive links for each task, while the y-axis shows performance shortfall, defined as 1 −AUROC using the held-out AUROC values reported in Table 2. Each point represents one prediction task; marker shape distinguishes the ncRNA source type, and color indicates the prediction context. Movement from the lower-left to the upper-right indicates increasing observed task difficulty, combining a larger candidate space with a larger AUROC shortfall. MET-related tasks occupy the lowest-difficulty region, Cancer-level tasks show intermediate difficulty, and CSMI-related tasks show the highest candidate-space burden and largest shortfall. The dashed horizontal line indicates the global model shortfall of 0.120.

### 3.4 Ablation Study Results

All three ablations reduced held-out AUROC relative to the full model. The largest reduction occurred under independent task training (−0.0416), followed by primary context backbone knockout (−0.0374) and message-topology randomization (−0.0259; Table 3).

**Table 3:** Held-out performance of the full model and selected ablation variants. Values are reported as mean ±standard deviation across ten independent runs.

| Model Variant | Accuracy | F1 Score | Precision | Recall | AUROC |
| --- | --- | --- | --- | --- | --- |
| Full model | 0.8206 ± 0.0109 | 0.8260 ± 0.0071 | 0.8032 ± 0.0257 | 0.8513 ± 0.0213 | 0.8801 ± 0.0118 |
| Independent task training | 0.7633 ± 0.0308 | 0.7839 ± 0.0166 | 0.7282 ± 0.0535 | 0.8557 ± 0.0519 | 0.8385 ± 0.0203 |
| Primary context backbone knockout | 0.7752 ± 0.0178 | 0.7919 ± 0.0155 | 0.7376 ± 0.0197 | 0.8551 ± 0.0188 | 0.8427 ± 0.0189 |
| Message-topology randomization | 0.7821 ± 0.0194 | 0.7978 ± 0.0119 | 0.7467 ± 0.0327 | 0.8583 ± 0.0245 | 0.8542 ± 0.0116 |

### 3.5 Case Study 1: External Disease-Level Assessment of Pancreatic Cancer miRNA Predictions

The top ten predicted miRNAs for pancreatic cancer (miR-137, miR-122, miR-128, miR-193b-5p, miR-381, miR-615-3p, miR-485-5p, miR-1297, miR-4709-3p, and miR-326) were cross-referenced against RNADisease v4.0 (Chen et al., 2023).

Nine candidates had pancreatic cancer records in RNADisease v4.0. No pancreatic cancer record for miR-4709-3p was identified in the prespecified database query, so it remained a model-prioritized candidate.

### 3.6 Case Study 2: External Literature Assessment of Model Predictions in Colorectal Cancer

For colorectal cancer, external literature assessment supported 6 of the top 15 miRNA–CSMI predictions and 8 of the top 15 circRNA–CSMI predictions with event-relevant experimental evidence (Tables 4 and 5). Predictions without matching event-level evidence in CRC were retained as prioritized candidates.

**Table 4:** External evidence status of top-ranked model-predicted miRNA–CSMI interactions in the colorectal cancer context.

| miRNA ID | metastatic event | Reference |
| --- | --- | --- |
| miR-326 | Cancer cell migration | (Wu et al., 2015) |
| miR-137 | Cancer cell proliferation | (Liu et al., 2023) |
| miR-1297 | Cancer cell migration | (Chen et al., 2014) |
| miR-572 | Cancer cell migration | (Wang et al., 2018) |
| miR-4500 | Cancer cell migration | (Yu et al., 2016) |
| miR-451a | Cancer cell migration | (Wu et al., 2021) |
| miR-3194-3p | Angiogenesis | No direct experimental evidence. |
| miR-3195 | Cancer cell migration | No direct experimental evidence. |
| miR-638 | Angiogenesis | No direct experimental evidence. |
| miR-330-3p | Angiogenesis | No direct experimental evidence. |
| miR-377 | Angiogenesis | No direct experimental evidence. |
| miR-105 | Angiogenesis | No direct experimental evidence. |
| miR-628-5p | Angiogenesis | No direct experimental evidence. |
| miR-4500 | Angiogenesis | No direct experimental evidence. |
| miR-940 | Angiogenesis | No direct experimental evidence. |

**Table 5:** External literature assessment of top-ranked model-predicted circRNA–CSMI interactions in colorectal cancer.

| circRNA ID | metastatic event | Reference |
| --- | --- | --- |
| hsa_circ_0006156 | Cancer cell migration | (Zeng et al., 2022) |
| hsa_circ_0001742 | Cancer cell proliferation | (Huang et al., 2024) |
| hsa_circ_0001178 | CRC to liver metastasis | (Ren et al., 2020) |
| hsa_circ_0006156 | Angiogenesis | (Zeng et al., 2020) |
| hsa_circ_0002360 | Cancer cell proliferation | (Chen et al., 2020) |
| hsa_circ_0005615 | Cancer cell proliferation | (Ma et al., 2020) |
| hsa_circ_0005615 | Cancer cell invasion | (Ma et al., 2020) |
| hsa_circ_0008234 | Cancer cell proliferation | (Wu et al., 2023) |
| hsa_circ_0002483 | Epithelial–mesenchymal transition | No direct experimental evidence. |
| hsa_circ_0000745 | CRC to lung metastasis | No direct experimental evidence. |
| hsa_circ_0003028 | Cancer cell migration | No direct experimental evidence. |
| hsa_circ_0001801 | CRC to liver metastasis | No direct experimental evidence. |
| hsa_circ_0000711 | Cancer cell migration | No direct experimental evidence. |
| hsa_circ_0006117 | Cancer cell migration | No direct experimental evidence. |
| hsa_circ_0004390 | CRC to liver metastasis | No direct experimental evidence. |

### 3.7 Case Study 3: Expression-Based Assessment of Predicted Migration-Associated miRNAs in Hepatocellular Carcinoma

To assess whether the model-prioritized HCC migration hypotheses showed independent expression support, the top 15 miRNA–CSMI predictions for the Cancer cell migration–Hepatocellular cancer context were compared with the external GSE67139 expression dataset. In this analysis, vascular invasion was used as an external metastasis-associated phenotype. VI-positive HCC tumors were compared with VI-negative HCC tumors.

Among the top fifteen predicted miRNAs, twelve were mappable in the GEO2R output, while three were not mapped to available probes. Ten of the mappable candidates showed adjusted-significant differential expression between VI-positive and VI-negative HCC tumors. The strongest expression-supported candidates included miR-200b, miR-29c, miR-143, miR-140, miR-34a, miR-455-3p, miR-19b-1-5p, miR-629-3p, miR-1273a, and miR-1272. miR-616-3p showed nominal significance but did not pass adjusted significance, while miR-137 was detected but not differentially expressed (Table 6).

**Table 6:**
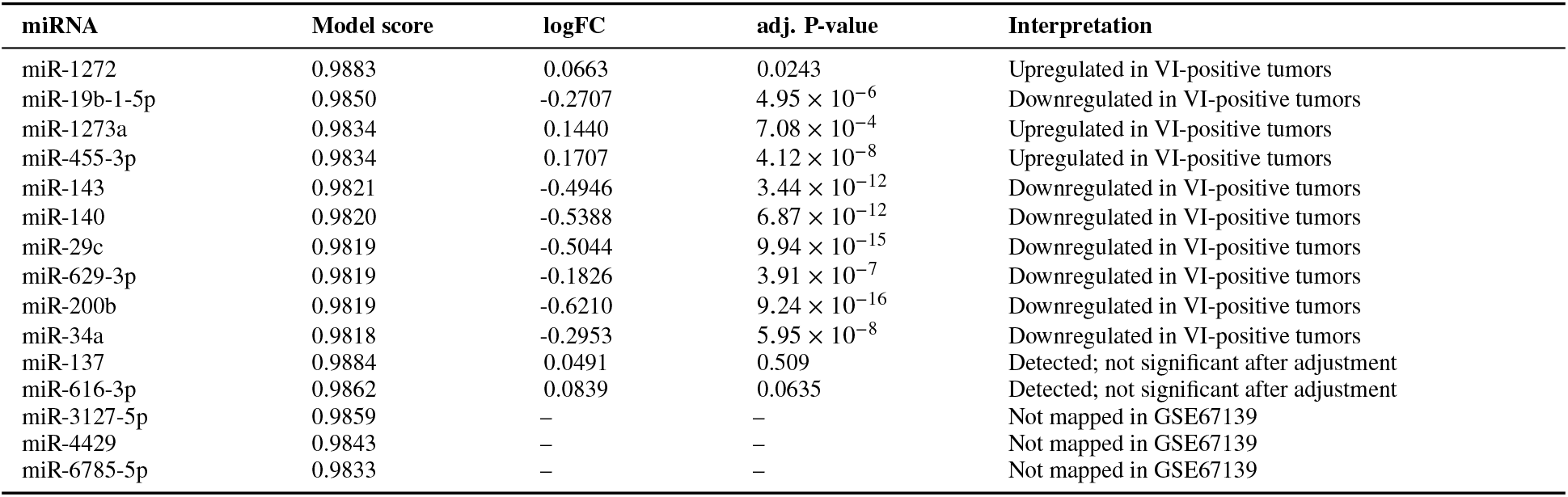
Expression-based assessment of the 15 top-ranked model-predicted migration-associated miRNAs in HCC using GSE67139. Positive logFC values indicate higher expression in vascular-invasion-positive HCC tumors. The table includes ten adjusted-significant candidates, three unmapped candidates, and two detected candidates that did not pass adjusted significance.

| miRNA | Model score | logFC | adj. P-value | Interpretation |
| --- | --- | --- | --- | --- |
| miR-1272 | 0.9883 | 0.0663 | 0.0243 | Upregulated in VI-positive tumors |
| miR-19b-1-5p | 0.9850 | -0.2707 | $4.95 \times 10^{-6}$ | Downregulated in VI-positive tumors |
| miR-1273a | 0.9834 | 0.1440 | $7.08 \times 10^{-4}$ | Upregulated in VI-positive tumors |
| miR-455-3p | 0.9834 | 0.1707 | $4.12 \times 10^{-8}$ | Upregulated in VI-positive tumors |
| miR-143 | 0.9821 | -0.4946 | $3.44 \times 10^{-12}$ | Downregulated in VI-positive tumors |
| miR-140 | 0.9820 | -0.5388 | $6.87 \times 10^{-12}$ | Downregulated in VI-positive tumors |
| miR-29c | 0.9819 | -0.5044 | $9.94 \times 10^{-15}$ | Downregulated in VI-positive tumors |
| miR-629-3p | 0.9819 | -0.1826 | $3.91 \times 10^{-7}$ | Downregulated in VI-positive tumors |
| miR-200b | 0.9819 | -0.6210 | $9.24 \times 10^{-16}$ | Downregulated in VI-positive tumors |
| miR-34a | 0.9818 | -0.2953 | $5.95 \times 10^{-8}$ | Downregulated in VI-positive tumors |
| miR-137 | 0.9884 | 0.0491 | 0.509 | Detected; not significant after adjustment |
| miR-616-3p | 0.9862 | 0.0839 | 0.0635 | Detected; not significant after adjustment |
| miR-3127-5p | 0.9859 | – | – | Not mapped in GSE67139 |
| miR-4429 | 0.9843 | – | – | Not mapped in GSE67139 |
| miR-6785-5p | 0.9833 | – | – | Not mapped in GSE67139 |

The hypothesis-generation and external evidence-assessment workflow for the three case studies is summarized in Figure 7.

**Figure 7:**
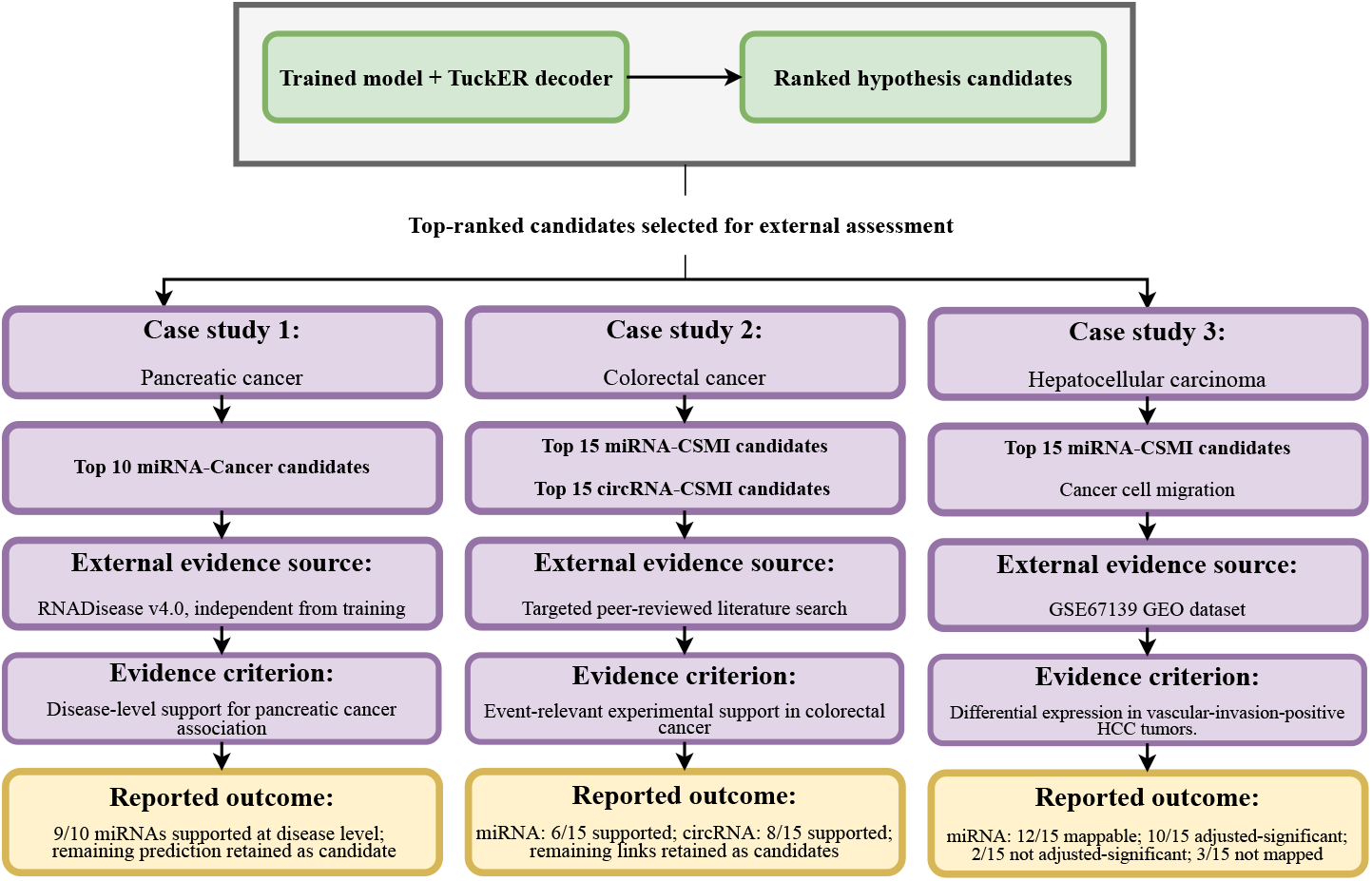
Decoder-guided hypothesis generation and external evidence assessment. Unobserved interactions were ranked as hypothesis candidates and assessed using external evidence.

### 3.8 High-Frequency Metastatic Contexts and Recurrent ncRNA Candidates

We examined predicted ncRNA–CSMI candidate associations across high-ranked cancer contexts to characterize the broader prediction landscape (Figure 8). Cancer cell migration in lung adenocarcinoma contained the largest number of retained candidate associations among the examined cancer–event contexts. Angiogenesis, cancer cell invasion, and cancer cell proliferation also contained substantial numbers of candidate associations in selected cancer types. Several circRNAs were repeatedly identified across the selected cancer contexts, including hsa_circ_0000745, hsa_circ_0000711, hsa_circ_0005615, and hsa_circ_0001801. Recurrent miRNA candidates included miR-3194-3p, miR-122-5p, miR-137, miR-572, and miR-1273a.

**Figure 8:**
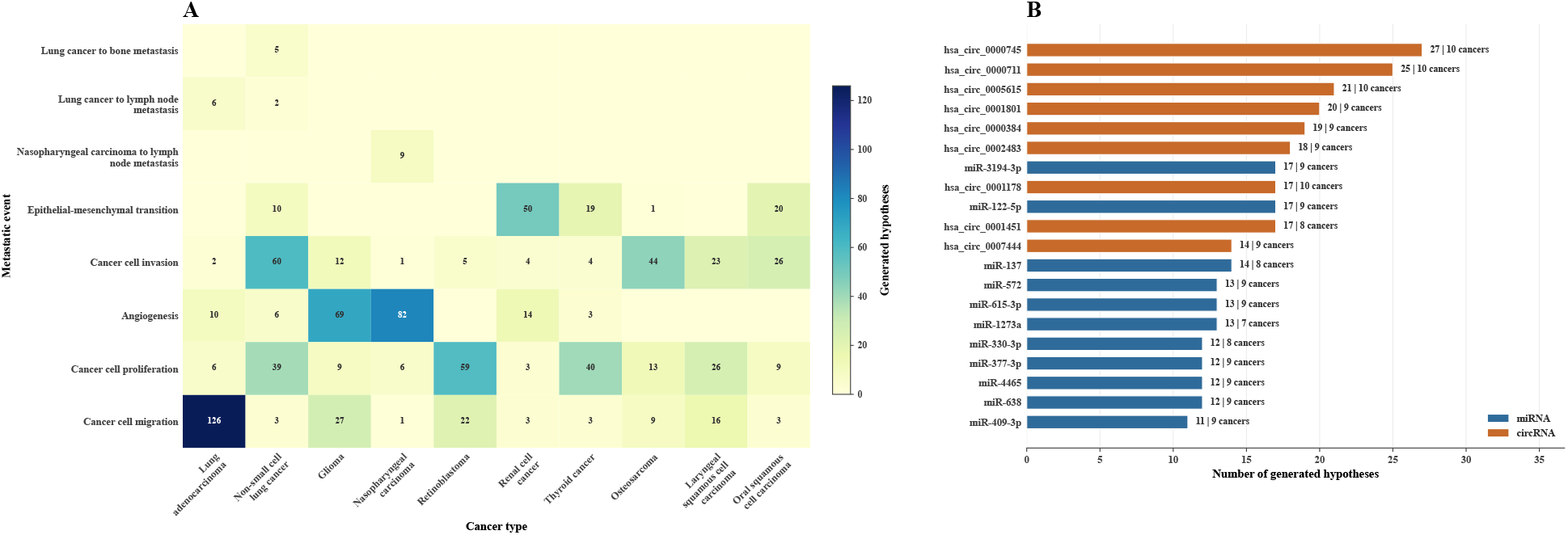
Global prediction landscape across high-ranked cancer contexts. (A) Heatmap of generated ncRNA–CSMI hypotheses across the ten highest-ranked cancer contexts. Columns represent cancer contexts, rows represent metastatic-event labels, and each cell reports the number of generated hypotheses for the corresponding cancer–event context. Darker colors indicate higher numbers of predictions. (B) Recurrent ncRNA candidate hubs across the same selected cancer contexts. Horizontal bars rank ncRNAs by the number of generated ncRNA–CSMI hypotheses in which they appeared. The label at the end of each bar indicates the total number of generated hypotheses for that ncRNA and the number of cancer contexts in which it was identified.

## 4 Discussion

Taken together, the task-wise results show that predictive performance decreases as the target context becomes more specific. MET-level tasks benefit from metastatic-event patterns shared across cancer types, whereas CSMI-level tasks require the model to resolve a specific cancer–event combination within a larger and sparser candidate space; Cancer-level tasks occupy an intermediate position between these two settings. CSMI prediction therefore represents the principal modeling bottleneck, while the strong MET-level performance shows that the shared representation captures robust event-level metastatic patterns.

Our hierarchical multi-task GNN extends ncRNA interaction prediction in cancer metastasis by jointly modeling miRNAs, circRNAs, cancer types, METs, and CSMIs. Unlike models focused on broad ncRNA–disease associations or pairwise interactions, including PDMDA (Yan et al., 2022), MVHGCN (Miao et al., 2025), BNPMDA (Chen et al., 2018), THGNCDA (Guo and Yi, 2024), DGNMDA (Lu et al., 2024), and circ2DGNN (Cen et al., 2024), our framework places predictions within both cancer-type and metastatic-event contexts, allowing the same ncRNA to be associated with different metastatic processes across cancers. It also differs from VGEA-LCME (Zhu et al., 2023), which uses variational graph autoencoders to predict lncRNA associations with metastatic events and organ-specific patterns, by focusing on miRNAs and circRNAs and jointly predicting six interaction types. Explicit CSMI nodes support cancer-specific and event-specific prediction, while expression-based features and learnable embeddings integrate information from multiple biological databases. Transformer Convolution, attention-based aggregation, and TuckER scoring provide multi-hop relational learning and multi-relational scoring across heterogeneous edge types. KS-CMI predicts circRNA–miRNA interactions without representing metastatic events (Wang et al., 2023), whereas graph triple-attention models operate on broad disease entities rather than explicit cancer–event contexts (Xuan et al., 2022). Our design choices reflect the context-dependent nature of ncRNA regulation in metastasis: the heterogeneous schema explicitly encodes cancer-type and metastatic-event information, allowing the encoder to learn cancer- and event-aware embeddings. These embeddings capture not only molecular identities but also contextual roles, including the context-dependent involvement of miRNAs in cancer invasion and metastasis (Baranwal and Alahari, 2010). Together, these design choices allow the model to represent biologically contextualized interactions instead of undifferentiated ncRNA–disease links.

The latent-space projections in Figure 9, generated using principal component analysis (PCA) (Pearson, 1901) followed by t-SNE (van der Maaten and Hinton, 2008), visualize the organization of the learned representations across heterogeneous biological entities and interaction contexts. In the node projection, the five node types occupy distinct regions of the shared latent space. In the edge projection, the six task-specific TuckER interaction-vector groups show relation-dependent organization within the decoder interaction space. The partial proximity among several edge groups indicates shared geometric structure across related ncRNA–Cancer and ncRNA–metastasis contexts.

**Figure 9:**
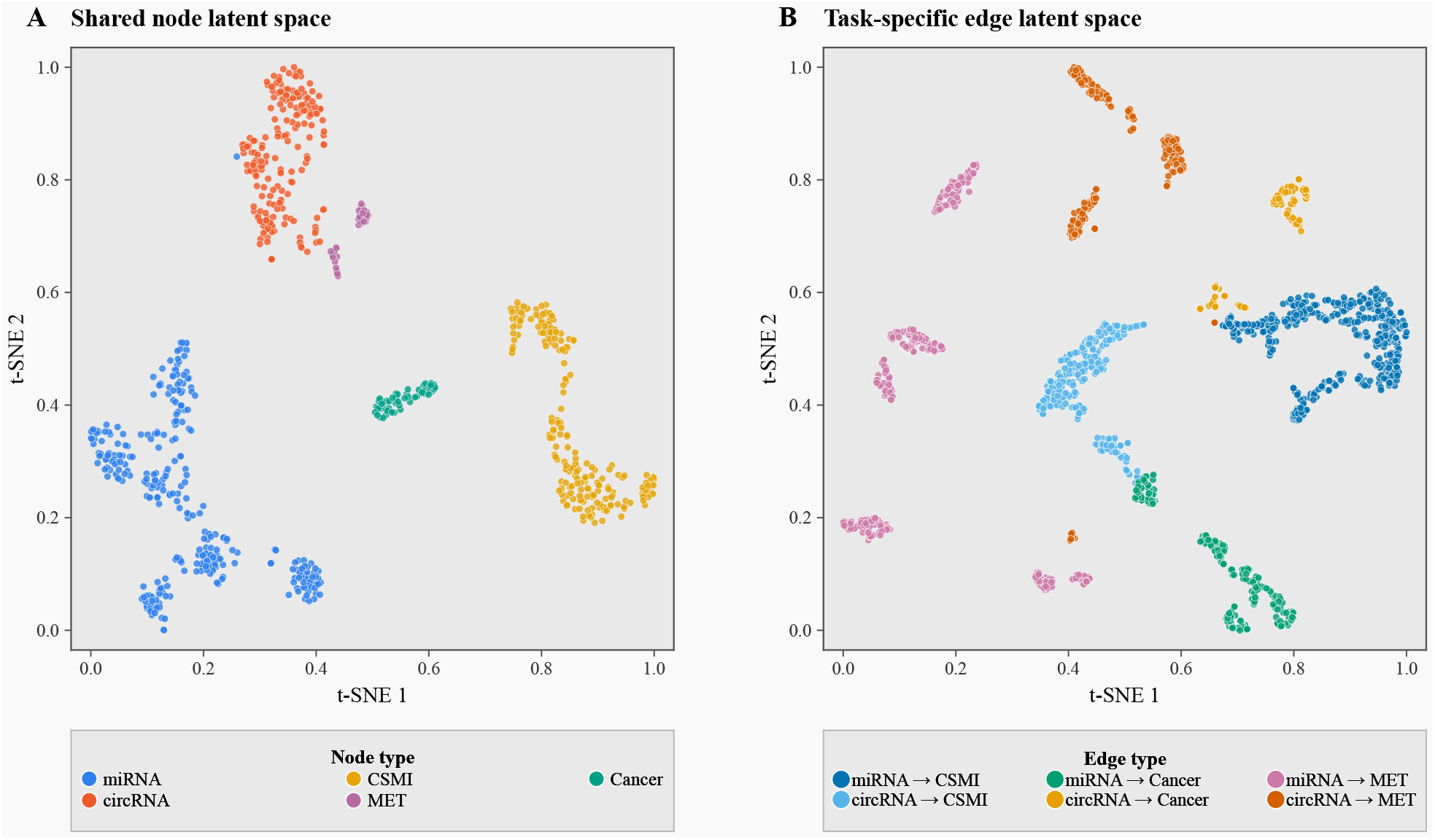
PCA–t-SNE visualization of the learned node and interaction representation spaces from the selected model checkpoint. (A) Two-dimensional projection of the final shared 12-dimensional node embeddings, grouped by node type. The node embeddings were standardized, PCA-transformed using all 12 components, and projected using t-SNE.(B) Two-dimensional projection of task-specific TuckER interaction vectors for observed positive edges from the six prediction tasks in the fitting message-passing graph, grouped by prediction task. For an edge (*h, r, t*), the interaction vector was defined as (**e**_*h*_**W**_*r*_) ⊙**e**_*t*_, where **e**_*h*_ and **e**_*t*_ are the task-adapted source and target embeddings and **W**_*r*_ is the relation-conditioned TuckER matrix. The edge vectors were standardized, independently PCA-transformed using all 12 components, and projected using t-SNE. Both projections used the same t-SNE settings, with PCA initialization and Euclidean distance. The resulting two-dimensional coordinates were rotated for display and independently min–max scaled to [0, 1].

The three external assessments evaluated complementary evidence levels: disease-level database records in pancreatic cancer, event-level experimental literature in colorectal cancer, and differential-expression evidence in vascular-invasion-stratified HCC tumors. Together with the global ncRNA–CSMI landscape, these analyses provide a prioritized set of candidates for targeted experimental testing.

## 5 Limitations and Future Directions

The reported metrics were computed on sampled positive–negative evaluation sets rather than on the full naturally imbalanced candidate space. They should therefore be interpreted as performance under the study’s controlled sampling protocol. In addition, the learning and evaluation pipeline treats unobserved pairs as negatives under a closed-world approximation. Because many unobserved ncRNA–context pairs are unlabeled rather than biologically absent, some sampled negatives may contain undiscovered positives. Model accuracy also remains constrained by the quality, completeness, and biases of public biological databases, which can introduce false negatives and context-specific noise. The evaluation is RNA-source-group-disjoint with respect to supervised target edges, but it remains transductive with respect to the complete node universe. Held-out RNA identifiers remain represented in the graph and can retain non-target relations and node-indexed parameters. The reported results therefore measure prediction of withheld target relations for RNA groups within a known heterogeneous graph, not inductive generalization to entirely new RNA entities. Cancer, MET, and CSMI nodes were present during fitting and were represented by trainable embeddings. The reported performance therefore applies to new RNA–context pairs involving known cancers and metastatic contexts and does not establish generalization to newly introduced cancer types, metastatic-event types, or CSMIs.

The hepatocellular cancer case study also uses vascular invasion as an external proxy for migration-related aggressiveness rather than as a direct measurement of cell migration. That validation therefore supports the metastatic relevance of the ranked candidates, but not migration-specific biology in a one-to-one sense.

Future work will address these limitations by expanding the graph with protein–RNA interactions, transcription factor binding data, and pathway databases, while integrating natural language processing for literature-mined evidence and RNA sequence features to generate richer node representations (Ivanisenko et al., 2024).

The “black-box” nature of deep GNN layers also limits interpretability. Future work may apply post-hoc explanation methods such as GNNExplainer to identify influential subgraphs and enhance model transparency (Ying et al., 2019). Finally, the current static graph cannot capture the dynamic rewiring of biological networks during disease progression and therapy. Future studies could explore dynamic and temporal GNN architectures using time-resolved data to model ncRNA network evolution and pinpoint drivers of metastatic transitions (Rossi et al., 2020).

## 6 Conclusion

We present a context-aware heterogeneous GNN that jointly models miRNAs, circRNAs, cancer types, metastatic events, and cancer-specific event contexts. RNA-group-disjoint testing and three external assessments showed that the model ranks ncRNA–context candidate associations across six prediction tasks. The framework provides a structured basis for prioritizing ncRNA–metastasis associations for experimental evaluation.

## 7 Data and Code Availability

The complete source code is publicly available on GitHub at https://github.com/Farzad-Midjani/ncRNA-Metastasis-Interactome-GNN. All datasets used in this study are publicly accessible through the databases cited in this paper.

## Notes

### Competing Interest Statement

The authors have declared no competing interest.

